# Methamphetamine-induced disruption of neuropeptide expression in mice

**DOI:** 10.64898/2026.06.20.733497

**Authors:** Nived Anidil Pathikkaran, Csenge Böröczky, Kyriaki Papageorgiou, Tomas Hökfelt, Tibor Harkany, Zsofia Hevesi

## Abstract

Psychoactive and psychotoxic drugs are particularly harmful, if their use coincides with critical developmental windows of brain maturation. Methamphetamine is one such stimulant with developmental exposure increasing seizure susceptibility and long-term neuronal maladaptation in children. Nevertheless, the extent at which infant and adult vulnerability to methamphetamine could differ in time-course and severity remains incompletely understood. Here, we developed a method to monitor methamphetamine-induced hyperactivity in infant mice at high temporal resolution, differentiate it from a biphasic response in adults, and link it to activity changes in cortical areas executing goal-directed (escape) behaviors in infant subjects when using *Fos* expression as a molecular surrogate. Subsequently, we hypothesized that methamphetamine could alter the expression and cellular distribution of ‘inhibitory’ neuropeptides, which, when co-released with fast neurotransmitters, could protect circuit plasticity by counteracting methamphetamine-induced hyperexcitability. Methamphetamine differentially altered somatostatin, cholecystokinin, and galanin expression in corticolimbic areas. These data suggest that methamphetamine can evoke age-specific neurocircuit modifications, at least in mice.

## Introduction

Clinical symptoms of acute exposure to methamphetamine, a potent psychostimulant, include hyperlocomotion, temporary cessation of food intake, increased core body temperature, and cardiovascular emergency^1,2,3^. In the brain, monoaminergic neurotransmission is particularly sensitive to methamphetamine action due to the drug’s ability to increase synaptic dopamine, serotonin, and norepinephrine levels^3,4,5^ by competing for neurotransmitter uptake at the plasmalemmal dopamine transporter (*Dat* /*Slc6a3*)^6^, and then reversing it to cause net efflux^7,8^. At the same time, methamphetamine disrupts the loading of neurotransmitter molecules into synaptic vesicles by vesicular monoamine transporter 2 (*Vmat2*/*Slc18a2*)^9^. Because methamphetamine increases neuronal activity (that is, action potential firing)^10,11^, the above mechanisms culminate into an overload of monoamines at (extra-)synaptic sites, and cause a generalized increase in network excitability^12,13,8^. It is therefore not inconceivable that the coincidence of methamphetamine exposure with critical periods of postnatal neurocircuit maturation (e.g., postnatal days (P)5-15)^14^ could be disruptive to neuronal wiring and function assignment^15^, affecting fundamental skills like rhythmic stepping and interlimb synchronization.

To reduce lasting maladaptation, the brain can rely on both dissipating excess neurotransmitters by fast enzymatic degradation and invoking intrinsic protective mechanisms. Neuropeptides present one distinct opportunity. Neuropeptides are short (<30-35 amino acids) bioactive peptides produced from prepropeptide precursors^16,17^, and co-released with fast neurotransmitters particularly upon repetitive neuronal activity (burst firing)^18,19,20,21^. Neuropeptides are not only numerous (>100 identified so far), but their temporal and spatial differences in availability and receptor heterogeneity suggest significant contributions to synaptic plasticity. By broad terms, neuropeptides can be subclassed as ‘excitatory’ *vs*. ‘inhibitory’^21,22,23,24^, with the physiological sign of action defined by their engagement of G_s_/G_q_ *vs*. G_i_ protein-coupled receptors (GPCRs), respectively. In the context of methamphetamine exposure, somatostatin (*Sst*)^25^, neuropeptide Y (*Npy*)^26,27^, and galanin (*Gal*)^28,29^ are of particular significance because of their known ability to reduce synaptic excitability^22,30,31^. Firstly, all are rapidly induced upon neuronal stimulation^16^. Secondly, their administration at pharmacological concentrations alters stimulus salience (increases reward), reduces stress/glucocorticoid-induced network hyperexcitability, and exerts antiepileptic and analgesic properties^32,33,34,35^. Thirdly, the expression of both the peptides themselves and their receptors is developmentally regulated^36^. Fourthly, experimental evidence places somatodendritic neuropeptide release to modulate neuritogenesis, axon guidance, and synapse selection in both cortical and subcortical neurocircuits during postnatal, activity-dependent maturation and refinement^37,38,39,40^. Thus, dysregulated neuropeptide signaling can be associated with neurodevelopmental disorders whether due to experiential factors or genetic predisposition^41,42,43,44^.

Nevertheless, if and to what extent neuropeptide systems are impacted by psychostimulants, when exposure coincides with neurocircuit maturation, remains incompletely addressed. Here, the behavioral sensitivity of infant (P8) and adult mice to methamphetamine was compared and then linked to age-specific changes in brain-wide neuronal activation, cellular stress, and neuropeptide expression. Corticolimbic areas were differentially affected, and showed the deregulation of ‘inhibitory’/protective neuropeptide components.

## Results

### Biphasic locomotor response to methamphetamine in adult mice

Methamphetamine (10 mg/kg)^10,45^ elicited an immediate increase in locomotion in adult mice (mixed for sex), consistent with its established psychostimulant properties^10,45^ (*F*_(2,69)_ = 40.31, *p* □0.0001; Fig.□1a). After a dormant period of ∼2 h when ambulation transiently returned to basal levels (but see below on stereotyped behaviors^1,46^), a secondary epoch of hyperactivity was observed (from 3 h on). Thus, the total distance moved in 4 h differed significantly between methamphetamine and vehicle-treated groups (*p* = 0.027; Fig. 1b). When using 5 mg/kg methamphetamine, a similarly biphasic locomotor response was found yet with higher activity scores retained in the trough of activity between 1 h and 3 h (Fig. 1a). The concentration-dependence of behavioral responses is consistent with high methamphetamine doses (>10 mg/kg) provoking stereotyped behaviors (e.g., excessive grooming)^1,46^, which could bias the automated detection of ambulation. Hyperactivity was accompanied by significantly reduced food seeking (*p* = 0.043; Fig. 1c) and elevated core body temperature (repeated-measures ANOVA: treatment: *F*_(1,7)_ = 12.09, *p* = 0.010; time: *F*_(2,14)_ = 11.91, *p* = 0.001; Fig. 1d), which persisted throughout the observation period. These data substantiate that methamphetamine elicits coincident behavioral and metabolic responses acutely in adult mice.

**Figure 1.**
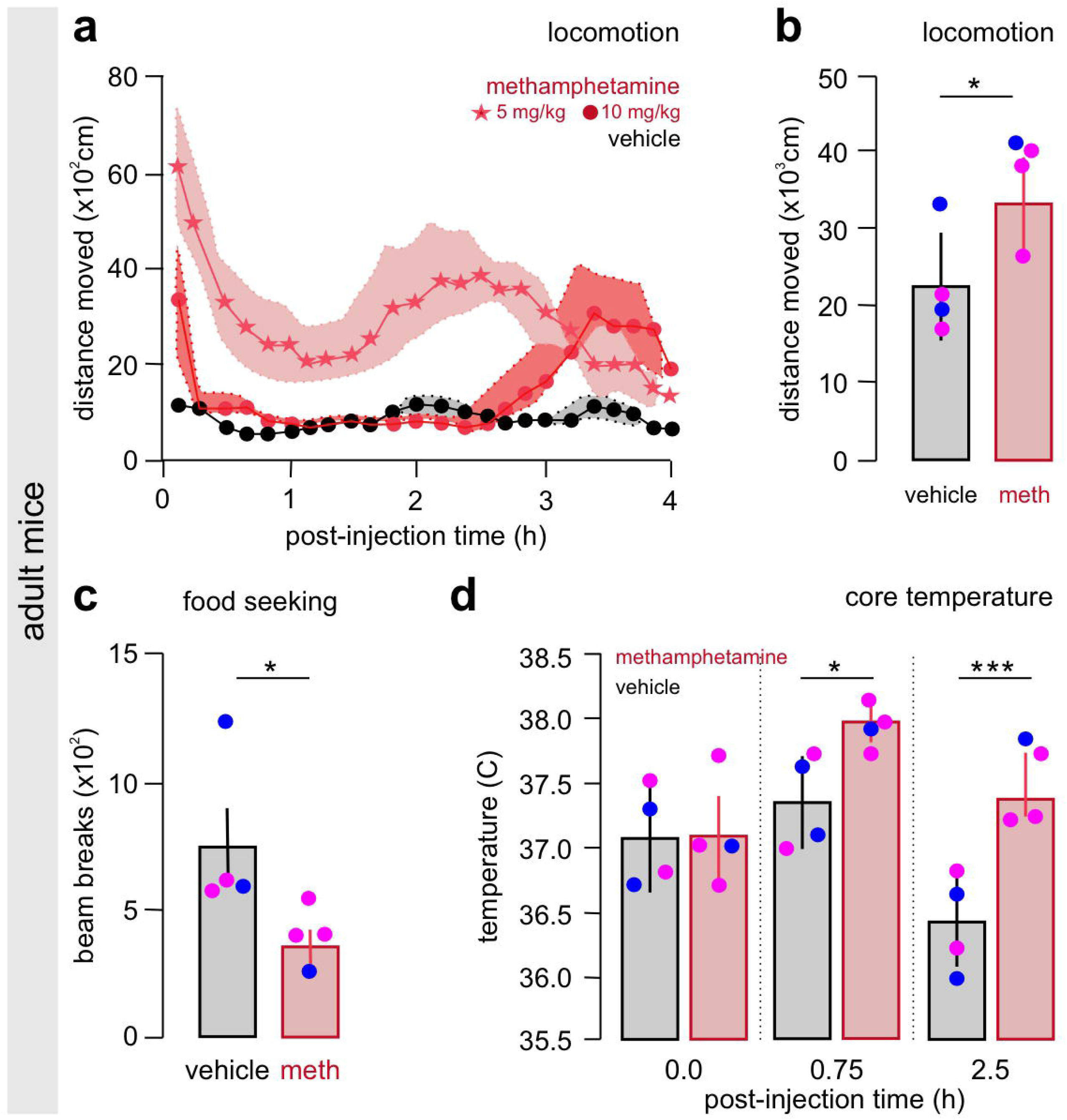
Metabolic parameters and biphasic hyperlocomotion following acute methamphetamine treatment in adult mice. (**a**) Continuous recording of ambulation for 4h revealed dose-dependent hyperlocomotion, characterized by acute (0-10 min; **b**) and delayed-onset (>2h) peaks (average values from mixed sex). (**c**) Methamphetamine (10 mg/kg, i.p.) suppressed food seeking (number of approach to the dispenser). (**d**) Simultaneously, methamphetamine increased core body temperature at both 45 min and 150 min (*n* = 4 mice/group). Data were expressed as means ± s.e.m., and statistically evaluated by one-way ANOVA (**a**), Student’s t-test (**b**, **c**) and two-way ANOVA (**d**). Solid circles show disaggregated data. Purple and blue data points denote female and male subjects, respectively. \**p*□< 0.05; \*\*\**p*□< 0.001.

### Behavioral effects of methamphetamine exposure in infant mice

Behavioral consequences of psychoactive drug exposure in infants are less known, because delayed eye opening (usually at postnatal day(P) 12-13)^47,48^ is seen as a formidable obstacle in rodent models. We sought to overcome this by establishing conditions for automated monitoring in a miniaturized open-field (9 cm in diameter; Fig. 2a). We used repetitive video recording for 5-min periods at 15 min, 2 h and 4 h after methamphetamine exposure. Pups were returned to the nest to avoid hypothermia and stress by excessive maternal deprivation^49,50,51,52,53^. Experiments were performed at ambient air temperature rather than thermoneutrality^54^ (∼29 °C) because a switch to higher temperatures could *per se* have biased mobility. On P8, methamphetamine (5□mg/kg, which is considered equipotent to 10 mg/kg in adult subjects)^55^ induced hyperactivity, which peaked at 15 min and then dissipated gradually (Mann-Whitney *U*-test: *p* < 0.0001 [15 min]; *p* = 0.014 [2 h]; *p* = 0.057 [4 h]; Fig. 2b,c). While vehicle-treated mice preferred the edge and walls of the arena, methamphetamine-treated infants did not exhibit patterned hyperactivity (Fig. 2c,d). These data show that methamphetamine-induced hyperactivity in infant mice persisted for hours.

**Figure 2.**
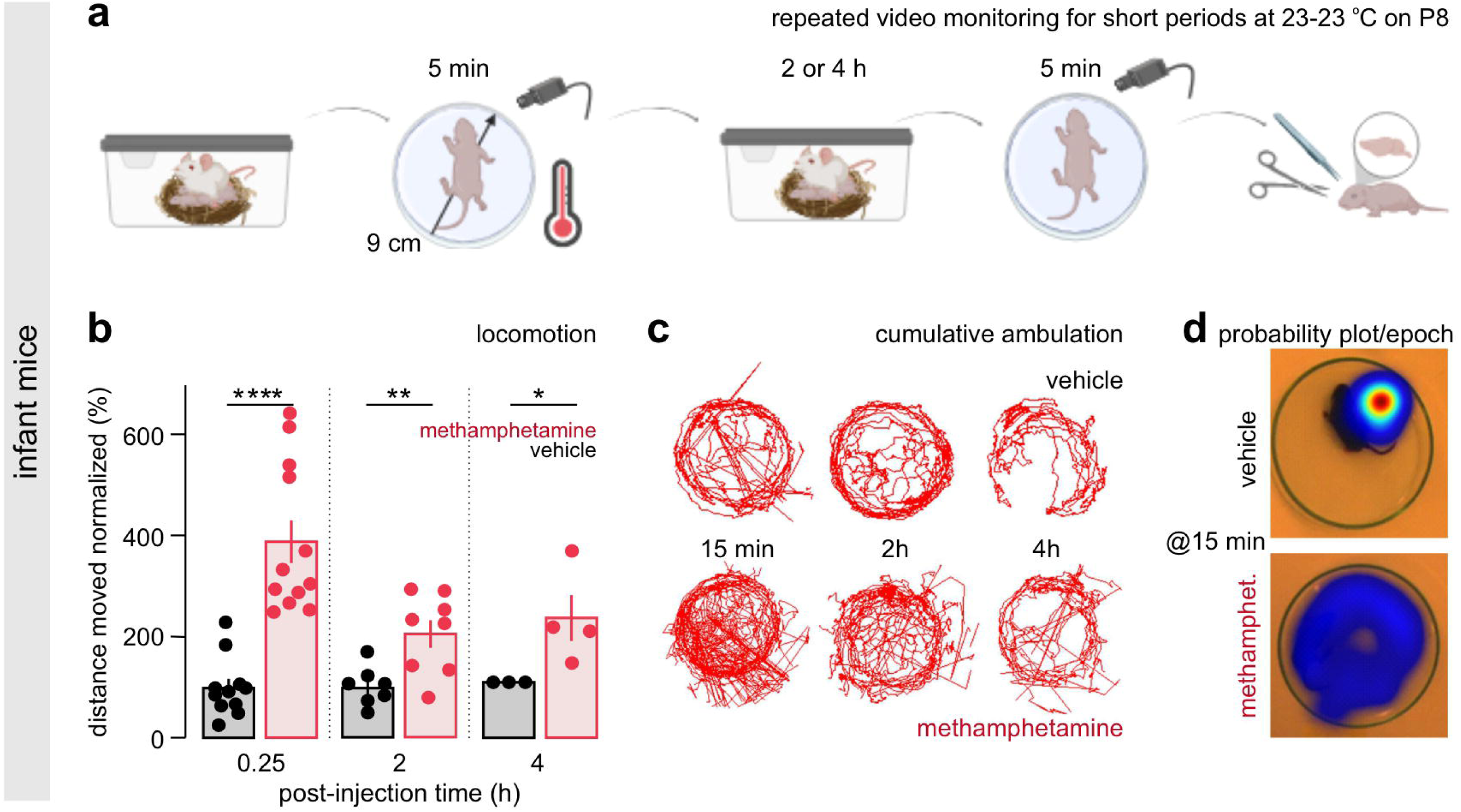
Methamphetamine effects on infant behavior. (**a**) Infant mice (P8) were first assessed in a miniaturized open-field for their locomotor activity upon methamphetamine exposure (5 mg/kg, *s.c.*). To prevent hypothermia, the pups were returned to the dam between consecutive video-recording sessions. After the second (2h) and third session (4h), pups were humanely euthanized and their brains used for neuroanatomy and biochemistry. Schematic representation was made in BioRender. (**b**) Methamphetamine (meth)-injected pups had continuous hyperlocomotion, with a peak response at 15 min (15 min: *n* = 11/12 vehicle/meth; 2h: *n* = 7/8 vehicle/meth; 4h: *n* = 3/4 vehicle/meth). (**c**) Representative activity maps to illustrate methamphetamine-induced hyperactivity. (**d**) Heat maps to confirm data in (**c**). Data were expressed as means ± s.e.m., and statistically evaluated by one-way ANOVA. Solid circles show disaggregated data. \**p*□= 0.05; \*\**p*□< 0.01; \*\*\*\**p* < 0.0001.

### Brain-wide neural activation upon methamphetamine exposure

Four hours after methamphetamine exposure, *Fos* mRNA measured by qPCR tended to increase in all brain areas of adult mice even if with considerable individual differences (*p* > 0.1; Fig. 3a)^56,10^. In eight-day-old mice, inter-subject variation was unexpectedly large regardless of the brain region examined (*p* > 0.1; Fig. 3b), precluding statistically justified differences. We considered these as natural variations in the temporal dynamics of immediate-early gene inactivation at the transcriptional level, which could be volatile after a pharmacological challenge^57^.

**Figure 3.**
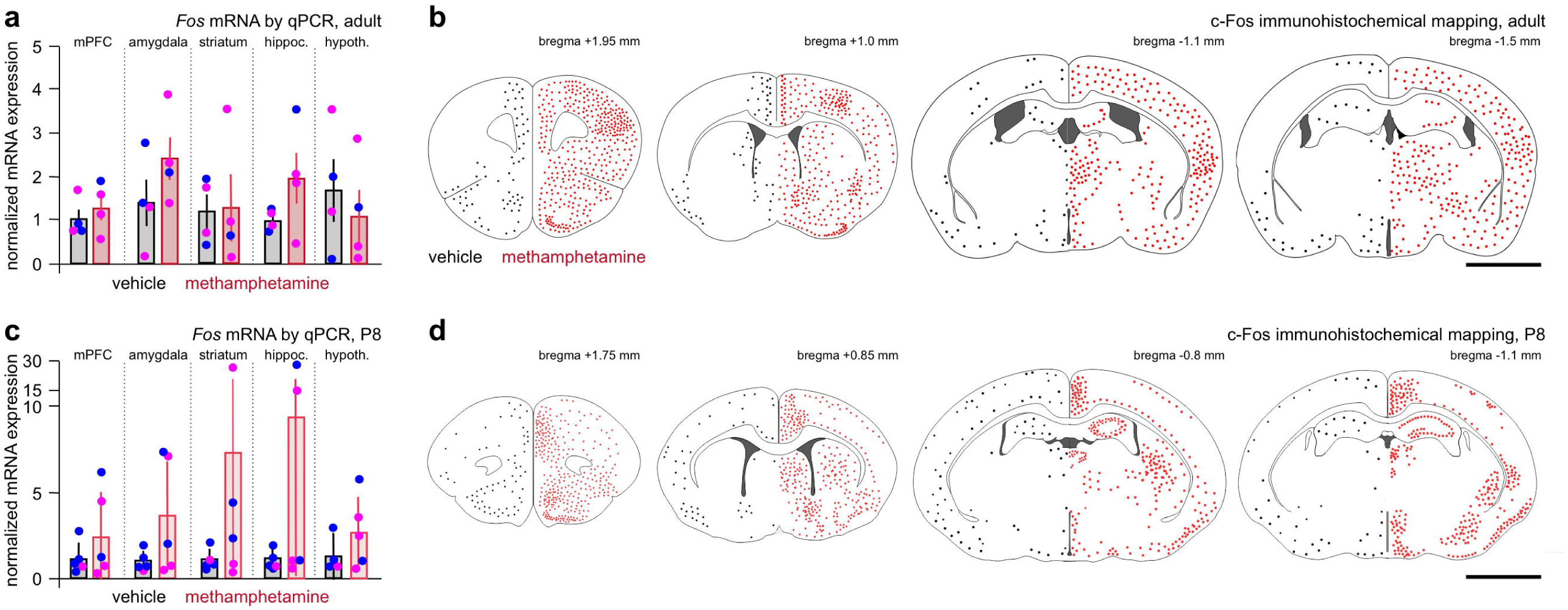
Brain-wide *Fos* activity mapping upon methamphetamine treatment. (**a**) In adult mice, methamphetamine had area-specific effects on *Fos* mRNA expression 2.5h post-injection. (**c**) c-Fos immunoreactivity significantly increased, including the prefrontal (mPFC) and somatosensory cortices, striatum, amygdaloid complex, thalamus and hypothalamus. Data represent the average from *n* = 3 animals. (**b**) In infant mice, *Fos* mRNA expression had unexpectedly large spread. No significant difference was found. (**d**) However, c-Fos immunoreactivity increased in all brain regions as in adults, with focal elevation in the retrosplenial and piriform cortices, striatum, habenula, and hypothalamus. Data per Bregma coordinate are the average labeling density from *n* = 3-4 animals. Data were expressed as means ± s.e.m., and statistically evaluated by one-way ANOVA. Solid circles show disaggregated data. Purple and blue data points denote female and male subjects, respectively. *Abbreviation*: P8, postnatal day 8.

Therefore, we carried out protein detection by forebrain-wide immunohistochemical mapping of c-Fos immunoreactivity in serial sections at both ages. In adult mice, c-Fos immunoreactivity dominated throughout the cortical mantle, with particularly dense cellular c-Fos immunolabelling in primary motor and somatosensory cortices. In addition, the nucleus accumbens, striatum, claustrum, habenula, hippocampus and hypothalamus, canonical target regions of psychostimulants^10,58,59^, were labelled (Fig. 3c). In P8 brains, c-Fos immunoreactivity concentrated in the retrosplenial and piriform cortices, hippocampus, hypothalamus, habenula, lateral septum, and striatum (Fig. 3d). Motor and sensory cortices exhibited sparse labelling. These findings suggest age-dependent c-Fos activation upon methamphetamine exposure, and are consistent with the activation of neural ensembles associated with spatial navigation, and reward in the infant brain^60,61^.

### Age-specific endoplasmic reticulum stress upon methamphetamine exposure

It is a common thought that younger brains encase higher levels of adaptive plasticity against adverse stimuli^62,63^. C/EBP homologous protein (CHOP, also termed GADD153) is a critical component of the apoptosis pathway triggered by endoplasmic reticulum stress^64,65^. CHOP downregulation, or decreased expression, is a mechanism for cell survival, particularly in cancer^66,67^. Conversely, CHOP upregulation is a sign of cellular vulnerability by affecting proteostasis^68^. Here, we explored if CHOP levels could support that methamphetamine, at the concentrations used, induces acute cell death at either age. In adult mice treated with 10 mg/kg methamphetamine, *Chop* mRNA levels were unchanged relative to vehicle-treated controls (*p* > 0.9 throughout; Fig. 4a). In contrast, markedly reduced *Chop* mRNA levels were determined at P8 that reached significance in the medial prefrontal cortex (mPFC; *p* = 0.056) and amygdala (*p* = 0.002, Fig. 4b). These data suggest an infant-specific adaptive response to methamphetamine likely aimed to enhance cell survival.

**Figure 4.**
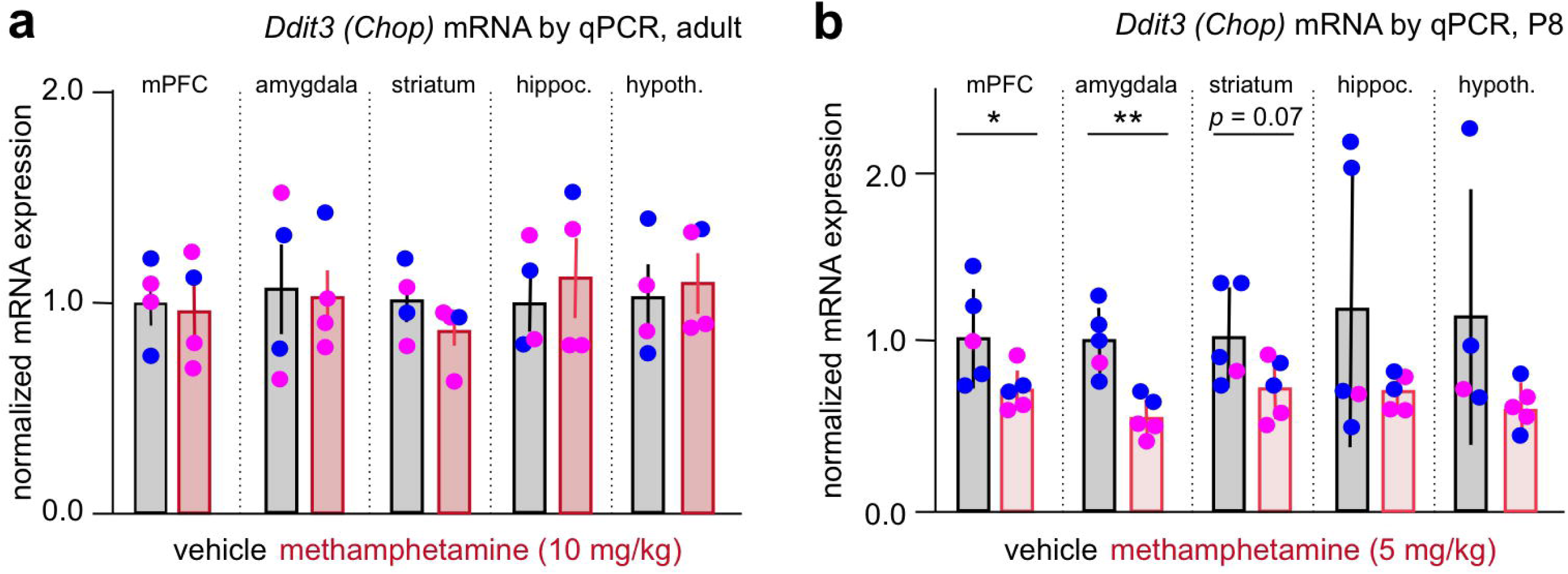
Methamphetamine does not induce endoplasmic reticulum stress. (**a**) In adult mice, methamphetamine did not affect mRNA levels for CCAAT-enhancer-binding protein homologous protein (*Chop*), which is rapidly induced by endoplasmic reticulum stress to mediate apoptosis (*n* = 4 mice/group). (**b**) In infant subjects, *Chop* mRNAs were in fact reduced by methamphetamine at 2.5h (n = 4-5 mice/group). Data were expressed as means ± s.e.m. and statistically evaluated by one-way ANOVA. Solid circles show disaggregated data. Purple and blue data points denote female and male subjects, respectively. \**p*□= 0.05; \*\**p*□< 0.01.

### Neuropeptide expression in methamphetamine-treated mice

Mechanistically, methamphetamine targets the mesocorticolimbic dopamine system to alter locomotion^4,5^, and even induces epilepsy upon repeated exposure^69^. Therefore, we have tested if expression of its molecular targets, *Slc6a3* (Dat1) and/or *Slc18a2* (Vmat2), is affected acutely in a region-specific fashion. Among the forebrain regions tested, both *Slc6a3* (*p* = 0.035; Fig. 5a) and *Slc18a2* (*p* = 0.066; Fig. 5b) mRNAs were downregulated in the striatum microdissected as soon as 2.5 h after methamphetamine exposure. These data validate the pharmacological model, and the concentration used.

**Figure 5.**
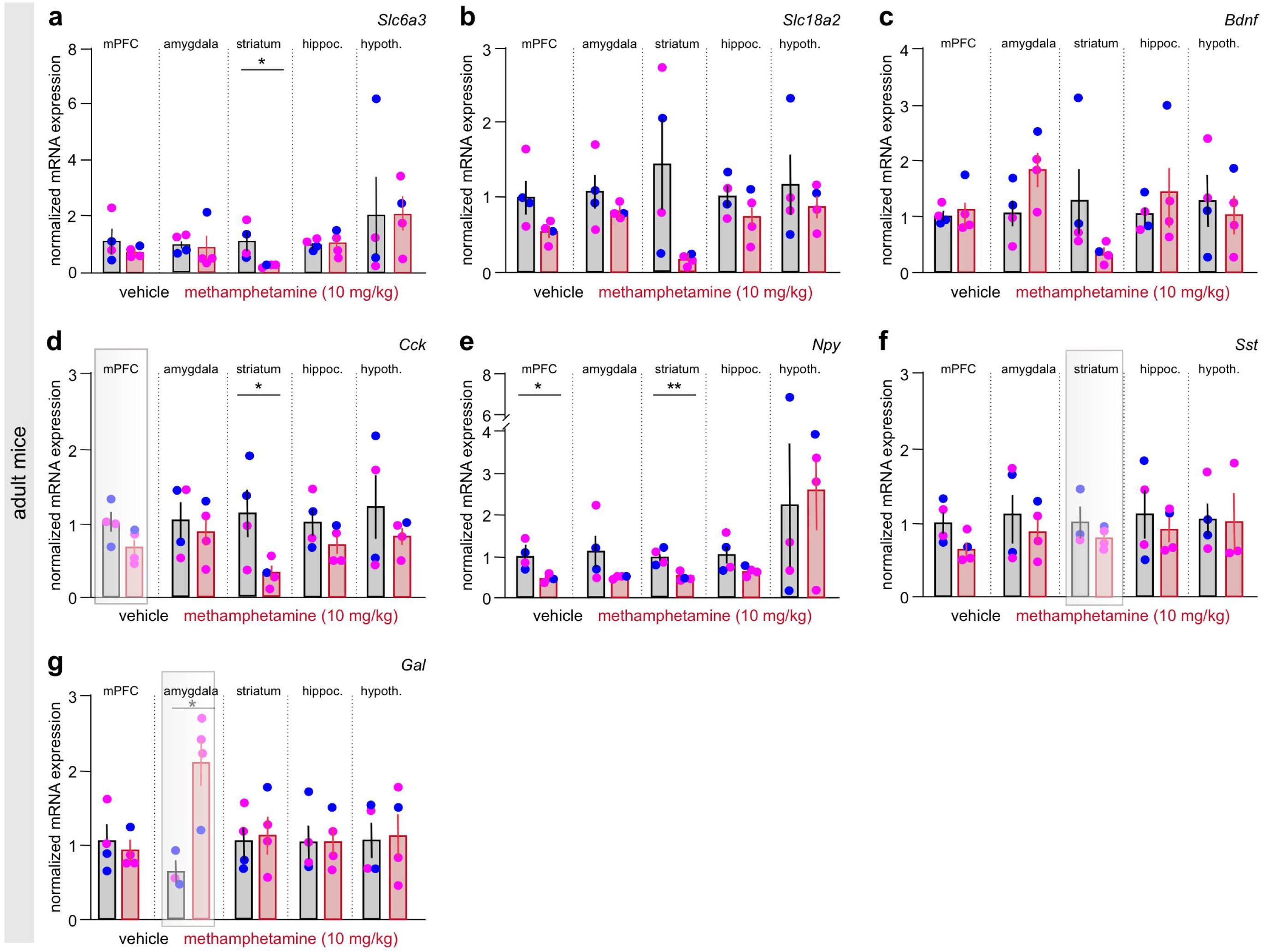
qPCR profiling of methamphetamine effects on neuropeptides in adult mice. (**d-g**) Methamphetamine reduced neuropeptide Y (*Npy*), somatostatin (*Sst*), and cholecystokinin (*Cck*) expression in prefrontal cortex (mPFC), amygdala, and striatum, whereas increased galanin (*Gal*) in the amygdaloid complex (*n* = 4 mice/group). Striatal downregulation of both *Slc18a2* (**b**) and *Slc6a3* (**a**) were taken as positive controls. Open rectangles (*gray*) indicate the neuropeptides and brain regions that had been assessed anatomically. Data were expressed as means ± s.e.m. and statistically evaluated by Students’s t-test. Solid circles show disaggregated data. Purple and blue data points denote female and male subjects, respectively. \**p*□< 0.05; \*\**p*□< 0.01.

Next, we assessed mRNA expression of neuropeptides considered either as morphogenetic cues during brain development (e.g., *Bdnf*, *Cck*)^70,71,72,73^ or neuroprotective in experimental disease models (e.g., *Gal*, *Npy*, *Sst* in epilepsy, neurodevelopmental disorders, and neurodegeneration)^74,75,76,77^. Methamphetamine did not induce consistent changes in *Bdnf* expression in adult subjects (Fig. 5c). Cck mRNA levels were reduced throughout, particularly in the striatum (*p* = 0.049; Fig. 5d). Likewise, *Npy* (mPFC: *p* = 0.017; striatum: *p* = 0.004; Fig. 5e) levels decreased. Changes in *Sst* mRNA content did not reach statistical significance (Fig. 5f). In contrast, *Gal* mRNA expression significantly increased in the amygdala (*p* = 0.016; Fig. 5g) but in no other area. These data suggest that neuropeptide expression rapidly changes presumably to reduce methamphetamine-induced neuronal hyperactivity.

At P8, *Slc6a3* mRNA levels were reduced in the mPFC (*p* = 0.048), and striatum (*p* = 0.051). Similarly, *Slc18a2* mRNA content decreased in the amygdala (*p* = 0.036), and even if to a lesser extent, in the striatum (*p* = 0.067; Fig. 6a,b), confirming the methamphetamine sensitivity of dopamine-rich areas.

**Figure 6.**
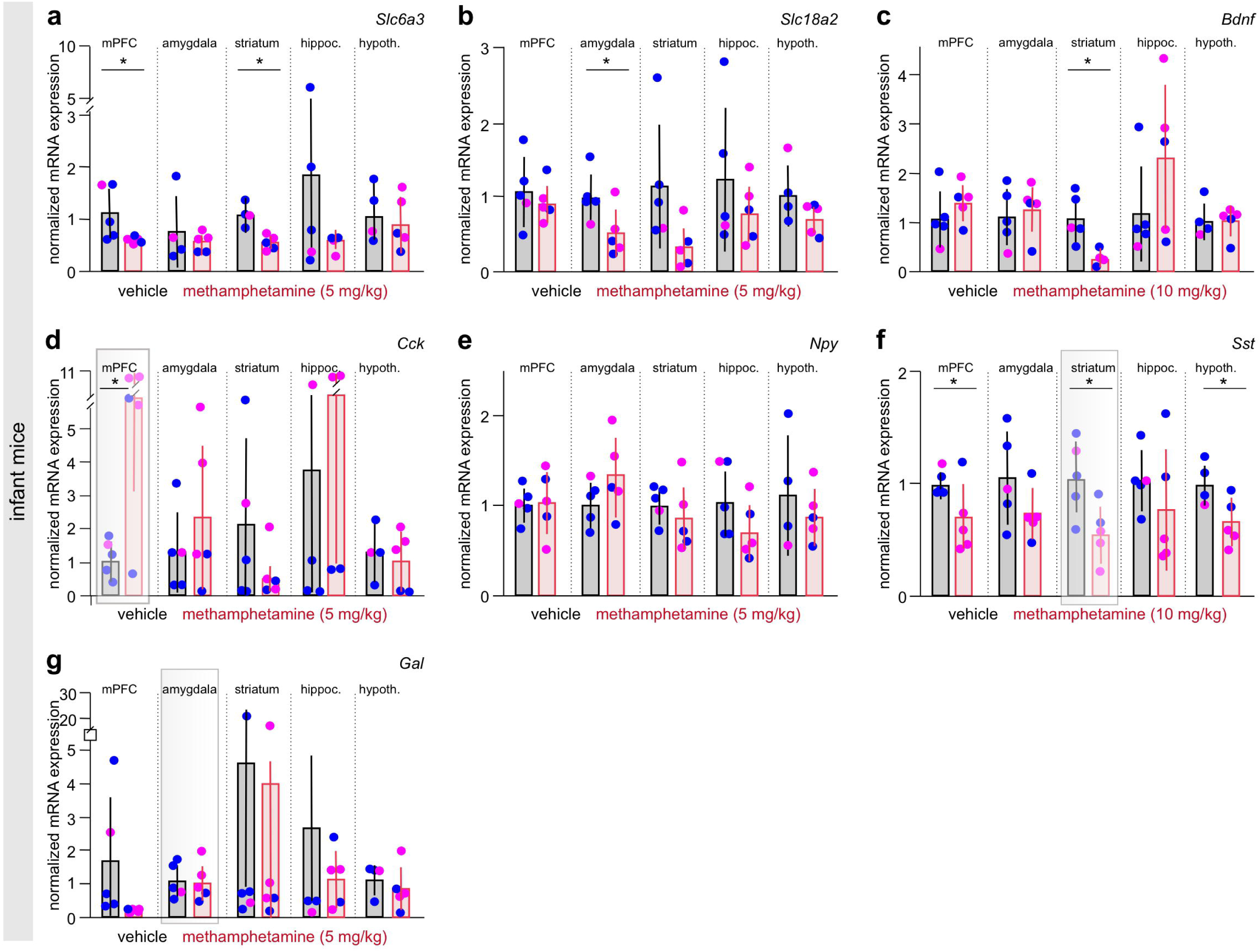
qPCR profiling of methamphetamine effects on neuropeptides in infant mice. (**d-g**) Methamphetamine reduced somatostatin (*Sst*) but increased cholecystokinin (*Cck*) expression in the striatum and prefrontal cortex (mPFC), respectively, at P8. Neither neuropeptide y (*Npy*) nor galanin (*Gal*) expression was affected (*n* = 4-5 mice/group). Striatal downregulation of both *Slc18a2* (**b**) and *Slc6a3* (**a**) served as positive controls. Open rectangles (*grey*) indicate the neuropeptides and brain regions that had been assessed anatomically. Data were expressed as means ± s.e.m. and statistically evaluated by Students’s t-test. \**p*□< 0.05. Solid circles show disaggregated data. Purple and blue data points denote female and male subjects, respectively.

*Bdnf* mRNA expression was significantly reduced in the striatum (*p* = 0.015; Fig. 6c). *Cck* mRNA levels were significantly increased in the mPFC (*p* = 0.017; Fig. 6d). *Npy* mRNA levels did not change (Fig. 6e). In contrast, *Sst* mRNA levels were reduced in the mPFC (*p* = 0.056), striatum (*p* = 0.0341), and hypothalamus (*p* = 0.036; Fig. 6f). *Gal* mRNA tended to be lower albeit without statistically significant differences (Fig. 6g). Cumulatively, these data suggest that methamphetamine broadly disrupts neuropeptide expression in the juvenile brain, particularly in areas that contribute to driving hypermobility.

### Neuropeptide mRNA localization

Bulk mRNA detection is insufficient to resolve cellular identity. We sought to overcome this limitation by cell-resolved fluorescence *in situ* hybridization (FISH), quantitatively comparing adult and P8 subjects. We performed a focused analysis on targets whose content differed, or tended to differ, upon qPCR profiling.

*Cck* mRNA expression levels differed in the mPFC of both adult and P8 mice, with opposing directionality of expression upon methamphetamine challenge. FISH validated these data by showing reduced *Cck*-positive(^+^) neuron numbers in adult subjects (Fig. 7a,a_1_), particularly in deep cortical layers. In contrast, the mPFC of P8 mice contained significantly higher numbers of *Cck*^+^ neurons (*p* = 0.044; Fig. 7d,d_1_), with methamphetamine-induced changes dominating in deep cortical layers.

**Figure 7.**
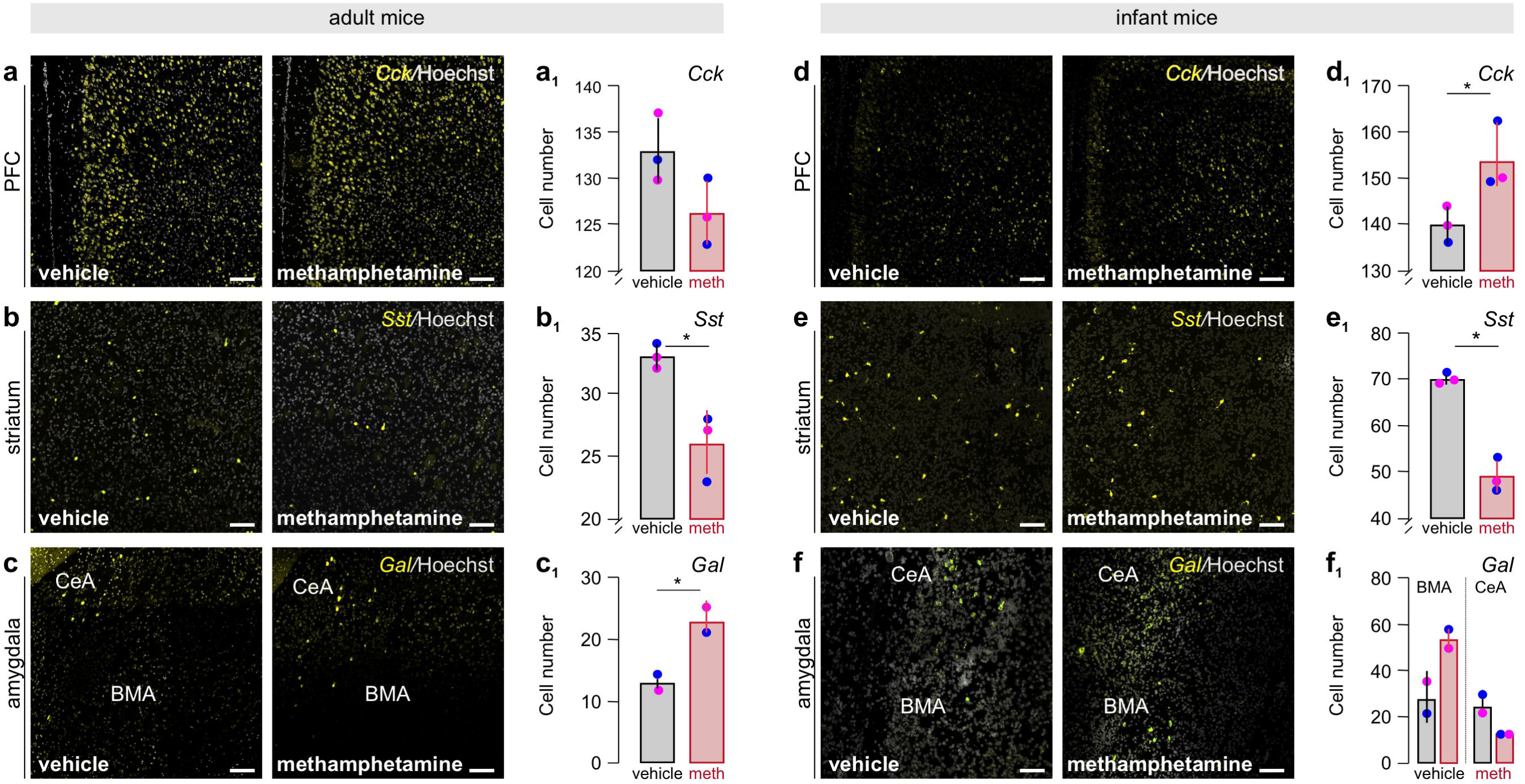
Cell-resolved, age-dependent effects of methamphetamine on some neuropeptides. Fluorescent *in situ* hybridization was used to anatomically justify altered neuropeptide expression upon methamphetamine exposure (as per Fig. 5,6). *Cck* expression was reduced in the medial prefrontal cortex (mPFC) of adults (**a,a_1_**), whereas increased in infants (**d,d_1_**). *Sst* mRNA significantly decreased in the striatum at both ages **(b,b_1_,e,e_1_**). *Gal* mRNA was increased in the amygdala at both ages (**c,c_1_**,**f,f_1_**). Note, that area-specific expression changes were observed at P8, characterized by a decrease in the central amygdala (CeA) and a concomitant increase in the basomedial subnucleus (BMA, **c_4_**). *Scale bars*: 100 µm (b_-_f), 200 µm (a,d). Data were expressed as means ± s.e.m. and statistically evaluated by Student’s t-test. \**p*□< 0.05; \*\*\**p*□< 0.001. Purple and blue data points denote female and male subjects, respectively. *Abbreviation*: CeA, central amygdaloid nucleus; L2/L5a/L6, cortical layers; BMA, basomedial amygdaloid nucleus; meth, methamphetamine; P8, postnatal day 8.

Next, we mapped *Sst* mRNA distribution in striata at both ages. Again, FISH corroborated the qPCR data by revealing reduced *Sst*^+^ cell numbers in both adults (*p* = 0.013; Fig 7b,b_1_) and infants (*p* < 0.001; Fig 7e,e_1_). These data showed that methamphetamine is a disruptor of striatal *Sst* expression, with its effect being independent of age.

Lastly, we determined *Gal* mRNA in the amygdaloid complex in both adulthood and at infancy, noting that qPCR did not bring about significant changes at P8. In adult mice, methamphetamine induced a significant increase in the number of cells expressing *Gal* mRNA in the central amygdaloid nucleus (CeA; *p* = 0.047; Fig. 7c,c_1_), supporting mRNA profiling in dissociated tissues. We then quantitatively assessed *Gal* mRNA at P8, and found a trend towards reduced cell numbers in the CeA (*p* = 0.079; Fig. 7f,f_1_). We noted that the basomedial amygdaloid nucleus (BMA) contained higher numbers of *Gal*^+^ neurons though (*p* = 0.083).

Overall, FISH not only confirmed qPCR data from bulk samples but allowed the differentiation of sub-regional changes in select neuropeptides, and their age-dependent modifications by acute exposure to methamphetamine.

### Immunohistochemical localization of galanin and somatostatin in methamphetamine-exposed brains

Neuropeptide mRNA expression is considered as rapidly inducible. Once processed, mature neuropeptides undergo rapid axonal transport for release^6,16,78,79^. Thus, neuropeptide accumulation in nerve endings can be used as either an anatomical feature to map brain areas or predict that neuropeptide inputs could, even if indirectly, build feed-forward expressional networks to amplify drug effects.

Firstly, we focused on galanin immunoreactivity in the CeA, which is known to receive galanin^+^ afferentation in mouse^80,75^. In adult subjects, methamphetamine upregulated galanin immunoreactivity in the CeA, which particularly concentrated around its c-Fos^+^ cell contingents (Fig. 8a). No such change in galanin content was seen in the BMA. In P8 brains exposed to methamphetamine, galanin input surrounded c-Fos^+^ cells predominantly in the BMA instead (Fig. 8c). These data infer differential accumulation of galanin in amygdaloid afferents at the two age groups studied.

**Figure 8.**
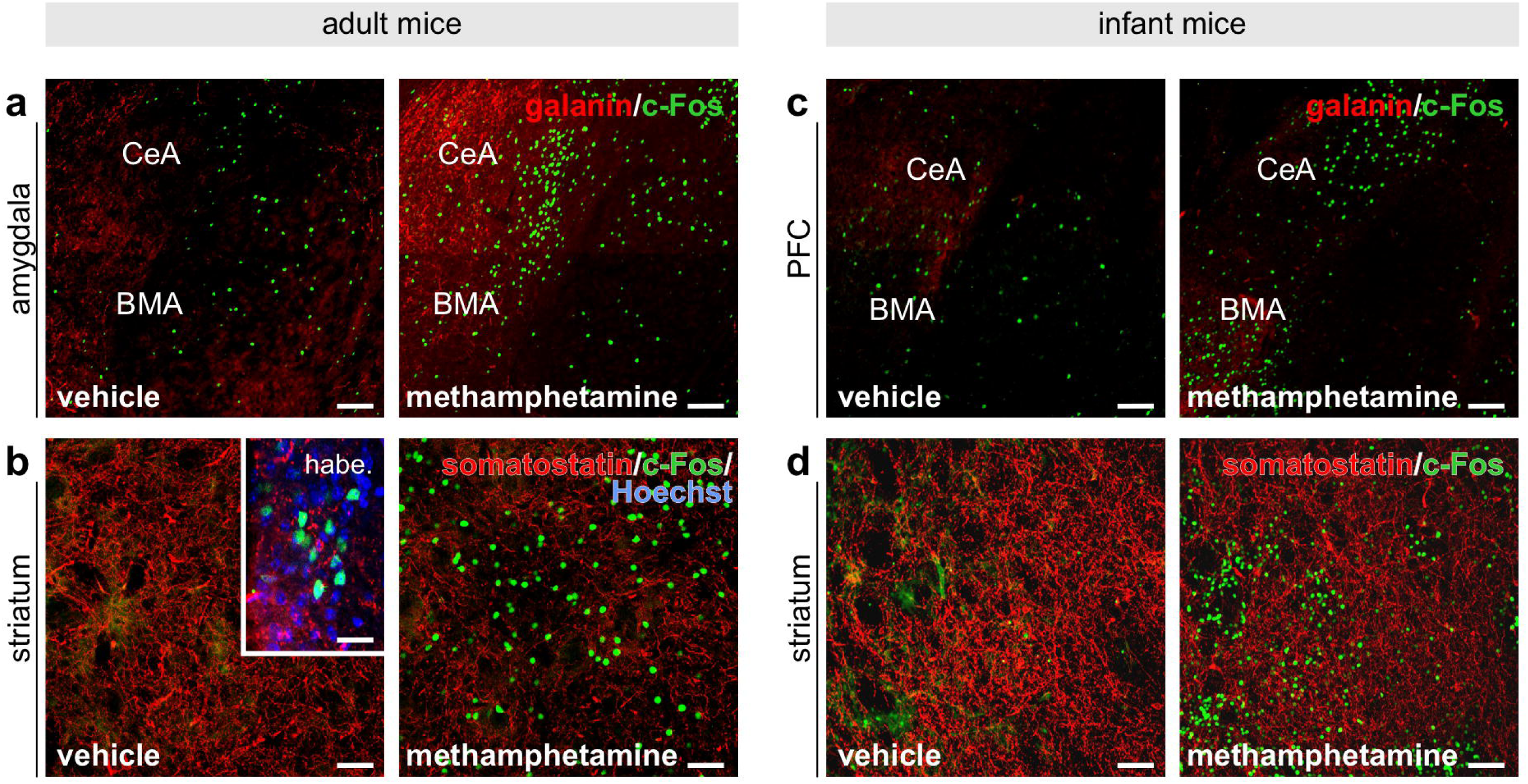
Immunohistochemical detection of neuropeptides upon methamphetamine exposure. (**a**) Galanin immunoreactivity was significantly increased in the amygdala of methamphetamine-exposed adult mice relative to controls. (**c**) No such change was seen in infant subjects, but area-specific expressional differences were detected. (**b**) The density of somatostatin-immunoreactive fibers was reduced in adults, and less so in infants (**d**) exposed to methamphetamine. c-Fos was used as positive control for methamphetamine-induced neuronal activation. (**b**, *inset*) Given the limited c-Fos immunoreactivity in adult controls, staining success was controlled by imaging the habenula, which contained many c-Fos^+^ neurons. *Scale bars*: 100 µm (a,c), 50 µm (b,d), 25 µm (b, inset). Data were expressed as means ± s.e.m. and statistically evaluated by Student’s t-test. \**p*□< 0.05; \*\*\**p*□< 0.001. *Abbreviation*: CeA, central amygdaloid nucleus; BMA, basomedial amygdaloid nucleus; meth, methamphetamine; P8, postnatal day 8.

Secondly, we mapped somatostatin immunoreactivity in the striatum at both ages, noting that besides intrinsic sources^81,82,83,84^, some somatostatin input to the striatum originates in pallidal structures^106^. Methamphetamine primarily increased c-Fos expression in striosomes at both ages (Fig. 8b,d), confirming treatment success. The density of somatostatin^+^ fibers in the striatal matrix was retained in methamphetamine-treated adult subjects (Fig. 8b), with somatostatin accumulating in ribbon-like structures, including putative nerve endings, around c-Fos^+^ cells at striosome-matrix interfaces. The density of somatostatin^+^ afferents did not qualitatively change at P8 (Fig 8.d). Notably, immunohistochemistry visualized many somatostatin^+^ neurons with bipolar or multipolar morphologies, some of which were c-Fos^+^. These data suggest that while somatostatin distribution becomes reorganized, even reduced, in the striatum of methamphetamine-exposed mice, a subset of striatal neurons can maintain somatostatin expression (Fig 8.b,d).

Taken together, these results suggest that acute methamphetamine exposure could induce the selective and regionally distinct reorganization of both intrinsic and afferent neuropeptide systems.

## Discussion

Here, we show that methamphetamine induces hyperactivity in both infant and adult mice with substantial difference in the time course of behavioral modifications. When using c-Fos as a surrogate of forebrain-wide neuronal activation^56,10^, we found that methamphetamine foremost affected motor and executive pathways at both ages, cortical activation, particularly in sensory areas, dominated in adult subjects. Excess neural activation was associated with changes in neuropeptide expression with NPY, somatostatin, cholecystokinin and galanin levels modified in age- and area-specific fashion. These findings underscore age-specific vulnerabiltiy of the brain to methamphetamine, at least in laboratory rodents.

The frequency of developmental toxin exposures, be these incidental, secondary (passive) or active intake, is increasing drastically^85,86,87^. On the one hand, the many recent societal trends with potential harm, e.g., recreational exposures or pressures to take party drugs, together with the many novel formulations that had become available (e.g., edibles, oils, drinks) increase the risk of adverse effects on brain function^88,89^. On the other hand, passive exposure to children (through medication during pregnancy, lactation, smoke inhalation or accidental ingestion) impinges on nervous system vulnerabilities to provoke lasting harm. While there is an increasing amount of literature on sex differences with boys considered more vulnerable than girls, consensus exists that children of both sexes carry considerable risk of brain maladaptation because drug exposure, even if for short periods, can change neurocircuit organization, thus potentially evoking long-lasting cognitive, emotional, or neuroendocrine/metabolic disruption^90,91,92^. Impaired neurocircuit functions can either manifest directly or constitute a ‘first subthreshold hit’ to sensitize the brain to later insults according to the ‘two hit hypothesis’ of neurodevelopmental disorders^93,94^. Our results focus on direct and acute effects of methamphetamine and show that this psychostimulant induces hour-long acute hypermotility in infants, with changes in general neural activity and neuropeptide levels. These data corroborate that early methamphetamine exposure carries major risk for neurological and psychiatric vulnerability^95,96^. Nevertheless, we acknowledge that our report is limited to acute time-points, with neither addressing long-term changes in neurotransmission within behavior-specific neurocircuits nor the molecular basis of altered neuropeptide content at the level of chromatin accessibility, transcription, or protein turnover. These questions offer exciting prospect for future research particularly when mechanistically comparing psychoactive drug profiles on developmental time-points, brain areas, and molecular targets.

Each organism resorts to an ever-changing repertoire of responses to environmental insults during its lifespan. Delayed eye opening in rodents determines the sequence of maturation for sensory systems^97,98,48,47^. It is therefore not unexpected that brain activation patterns would differ between blind infants and adults. Indeed, preferential activation of neural systems underpinning representation and navigation in, and reactions to complex environments, such as the retrosplenial cortex and mPFC for spatial context and decision making, respectively, were dominant in infants on P8^99^. In contrast, brain areas shaping context-dependent, goal-directed behaviors merging sensory input, spatial memory, and executive control to guide action were affected by methamphetamine in adults^100^. At the behavioral level, infants are likely limited to resort to stereotyped behaviors (e.g., grooming, saliva spreading/licking) that adults use as defenses to limit hyperthermia^1,46^. Thus, age-dependent behavioral response repertoires to methamphetamine are rooted deep in how experience (or the lack of) defines neurocircuit organization and sensitivity.

Acute methamphetamine exposure elicited coordinated changes in the expression patterns of galanin, cholecystokinin (CCK), and somatostatin (SST) at both transcript and protein levels in adult and P8 mice, pointing to a rapid engagement of neuropeptidergic signaling systems across developmental stages. Given that these neuropeptides critically regulate synaptic gain, inter-neuronal communication, and circuit-level excitability, their dynamic modulation is likely to reflect an intrinsic homeostatic response to psychostimulant-induced network perturbation. In particular, galanin is positioned to constrain excessive monoaminergic activity through inhibitory neuromodulatory actions^101,102^, whereas CCK^103^ and SST^104^ define functionally distinct interneuron populations governing oscillatory synchronization and inhibitory tone. The parallel responsiveness of these neuropeptide systems in the immature brain is especially notable, as it suggests that methamphetamine-sensitive signaling modules are already operational during early postnatal circuit assembly. Thus, neuropeptide remodeling may represent a conserved adaptive mechanism through which neuronal networks transiently recalibrate their excitatory–inhibitory balance following acute pharmacological challenge.

On a methodological note, assessing infant behaviors in rodents is challenging because the choice of tests is limited to assess subjects before eye opening. Geotaxis, tail hanging assess responsivity to passive stressors^105^. Here, we miniaturized an open-field to determine motor coordination and ambulation noting that at P8 whisker activity is likely sufficient to help infant mice to circumnavigate a circular arena with walls high enough to present boundaries^37^. At the same time, 5-min epochs were used to reduce the risk of hypothermia. Even more, short periods of observations likely prevented stress associated with maternal deprivation, a condition established by separating pups from the mother repetitively between P2-P14 for more than 120-180 min/day^52,53^. Thus, data provided in this report could genuinely capture methamphetamine-induced hyperactivity rather than a desperate attempt by the pups to find their dams and nest.

Together, our findings identify galanin, CCK, and SST signaling as conserved neuropeptidergic substrates engaged by acute methamphetamine exposure across development. The divergent behavioral dynamics in infant and adult mice further suggest age-dependent adaptations in neuronal network homeostasis.

## Methods

### Animals, husbandry, ethical considerations and drug treatment

C57BL/6J animals (Janvier) of both sexes were used throughout (adult: *n* = 9 females and 7 males, P8: *n* = 23 pups from 3 damns, mixed for sex). Data were disaggregated for sex where relevant. Experiments on live animals conformed to the 2010/63/EU directive and were approved by regional ethical authorities (Jordbruksverket/Stockholms djurförsöksetiska nämnd, Sweden: 221-2025). Animals were housed conventionally at 55% humidity and a 12-h/12-h light cycle. Mice had *ad libitum* access to food and water. Care was taken to minimize the number and the suffering of the mice used.

Methamphetamine (Sigma) was dissolved in physiological saline solution at a final concentration of either 5 mg/kg or 10 mg/kg freshly each day. Drug concentrations were corrected for bodyweight and given at a 6 μl/g volume. Equivolumetric physiological saline solution was used as vehicle throughout. Pups and adults received *s.c*. and *i.p*. injections, respectively.

### Behavioral assessment

Adult mice (*n* = 4 mice/group, 10-week-old) were placed individually in PhenoTyper cages (30 cm in diameter; Noldus) within incubators (Tecniplast) maintained at 23 °C. Following a two-day habituation period, animals received either 10 mg/kg methamphetamine or saline injections with their locomotion and food seeking continuously monitored during a 4-h period. Body temperature was measured at 15 min, 2 h, and 4 h by a rectal probe (BAT-10 Thermometer, Physitemp). Behavioral data were analyzed in EthoVision XT17 (Noldus).

To test if methamphetamine induced hypermobility in infant mice (*n* = 12/group for 15 min, *n* = 8/group for 2 h, *n* = 3-4/group for 4 h; mixed for sex throughout), pups on P8 were randomly assigned to the treatment groups while powering the experiment with both biological and technical replicates for both sex. Behaviors were assessed by placing the pups in glass Petri dishes (9 cm in diameter) on non-reflective backgrounds for improved signal-to-noise ratio and precise video tracking. Their motility was assessed in 5-min epochs at 15 min, 2 h and 4 h post-injection at 22-23 °C. Pups were returned to their respective nests to minimize stress and prevent hypothermia between recording sessions. Behavioral data were analyzed using EthoVision XT17 (Noldus), with track plots and heat maps generated from the recorded data.

### qPCR

Immediately after the behavioral tests, mice were terminally anesthetized with isoflurane (5%, flow rate: 1 L/min), decapitated, and their brains freshly dissected out for either mRNA extraction or fluorescence *in situ* hybridization (*see below*). Quantitative PCR (qPCR) reactions were performed on microdissected brain samples (*n* = 3-5/group/age, both sexes). Total RNA was extracted using an RNA Mini Kit (Bio-Rad) with its concentration determined on a NanoDrop. RNA samples (10 ng/μl, total input 200 ng/reaction) were reverse transcribed using a high-capacity cDNA reverse transcription kit (Thermo Fisher). qPCR reactions were performed after an initial denaturation step at 95 °C for 3 min followed by 35 cycles of 95 °C for 1 min denaturation, annealing and extension at 60 °C for 1 min each, and dissociation at 72 °C for 2 min on a CFX96 real-time PCR (Bio-Rad). Primer pairs were designed to amplify short fragments of *Fos, Gal, Cck, Sst, Npy, Bdnf, Slc6a3, Slc18a2* and *Chop* genes for mice (Table 1). TATA-binding protein (*Tbp*) served as internal control. For each gene, the respective melting curve showed a single product peak with no trace of primer dimers or other artifacts. Data were analyzed by using CFX Maestro v.5.3.022.1030 (Bio-Rad).

**Table 1.**
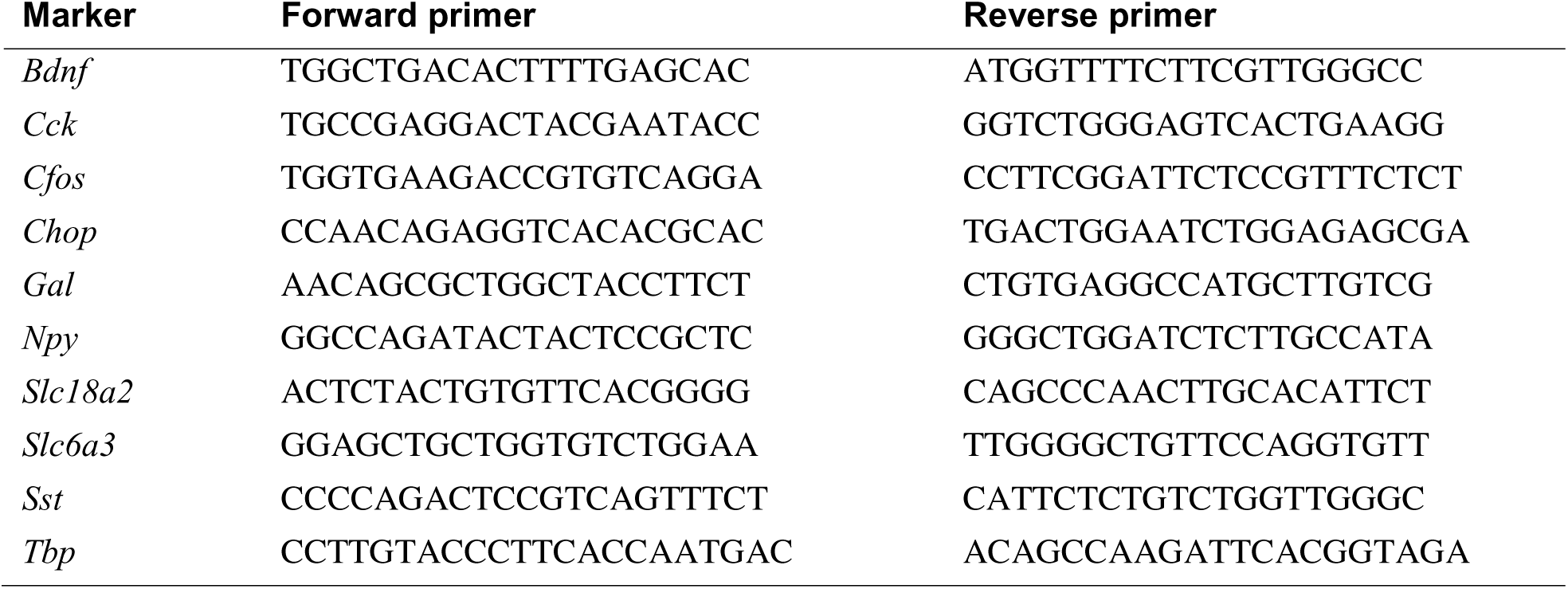
Primer sequences used in this study. qPCR reactions were performed with primer pairs amplifying short fragments for each gene. Primer pairs were custom designed to efficiently anneal to homologous nucleotide sequences in mouse.

### Fluorescence *in situ* hybridization (FISH)

Brains of both ages were embedded in optimal cutting temperature compound (OCT; Sakura) in plastic molds, flash-frozen on dry ice, cryosectioned at a thickness of 14 µm on a Leica CM1860 cryostat microtome and serially mounted onto SuperFrost^+^ glass slides (Thermo Fisher). Sections were immersion fixed in 4% paraformaldehyde (PFA) in 0.1M phosphate buffer (PB, pH 7.4) at 4 °C for 15 min, rinsed in 0.1M PB, and dehydrated through an ascending ethanol gradient (50%, 75%, and 100%; 5 min each). Multiplex *in situ* hybridization was performed using the HCR v3.0 protocol (Molecular Instruments) with probe sets for *Gal, Cck*, and *Sst*. Following hybridization, sections were counterstained with Hoechst 33,342 (Sigma; nuclear stain), air-dried, and coverslipped with Entellan (Sigma).

### Immunohistochemistry

Mice of both ages were transcardially perfused with 4% paraformaldehyde (PFA) in 0.1M phosphate buffer (PB, pH 7.4) using a 50 ml syringe. Brains were removed, post-fixed at 4 °C overnight, and cryoprotected in 30% sucrose in 0.1M PB. Brains were cryosectioned at 50-µm thickness (Leica CM1860), and processed as free-floating sections by first incubating them in 0.1M PB containing 5% normal donkey serum (NDS; Jackson ImmunoResearch) and 0.3 % Triton X-100 (Sigma) at 22-24 °C for 2 h. Subsequently, tissues were exposed to combinations of primary antibodies: guinea pig anti-cFOS (1:1,000; Synaptic Systems), rabbit anti-galanin (1:2,000; Peninsula), and mouse anti-somatostatin (1:500; GeneTex), diluted in 0.1M PB also containing 0.1% NDS and 0.3% Triton X-100 at 4 °C for 72 h. After repeated rinses in 0.1M PB, sections were incubated with species-specific secondary antibodies conjugated to carbocyanine (Cy2, Cy3, or Cy5; raised in donkey, 1:500; Jackson ImmunoResearch) at 4 °C overnight. Hoechst 33,342 (1:10.000; Sigma) was routinely used as nuclear counterstain. After rinsing in 0.1M PB, sections were dipped in distilled water, mounted on fluorescence-free glass slides, air-dried, and coverslipped with Entellan (Sigma).

### Imaging and quantification

Sections processed for either FISH or immunohistochemistry were imaged on an LSM880 confocal microscope (Zeiss) with the pinhole set to 1 airy unit when using a 20×/0.6 NA objective. Images representative of data from *n* = 2–3 mice/group were included in the figure panels. Multi-panel images were assembled in CorelDraw (v. 2022, Corel Corp.). Quantification was performed in ZEN 2012 (Blue Edition, Zeiss). For *Cck* and *Gal*, a standardized area of 0.1114 mm² per section was analyzed, whereas for *Sst* a larger area of 1.9 mm² was selected. The total number of cells within each region was counted, with the treatment effect computed at either age.

### Statistics

Data were expressed as means ± s.e.m. throughout. Data per treatment group were first evaluated for normality using the Shapiro-Wilk test. If passed, further data processing was based on parametric tests, including one-way ANOVA, Student’s *t*-test (two-tailed, unpaired, group design), or repeated-measures ANOVA (Fig. 1d). Alternatively, the Mann-Whitney *U*-test (rank sum) was employed in Fig. 2. Statistical analysis was performed in GraphPad Prism 8. A *P* value < 0.05 was considered statistically significant.

## Acknowledgements

The authors thank Dr. V. Cinquina for advice on CHOP analysis, and L. Glatt for her expert technical assistance. This work was supported by the Austrian Science Fund (10.55776/COE16; T.Ha.), the Swedish Research Council (2023-03058; T.Ha.), Novo Nordisk Foundation (NNF23OC0084476; T.Ha.); and the European Research Council (FOODFORLIFE, 2020-AdG-101021016; T.Ha).

## Author contributions

T.Ha. and Z.H. conceived the project. T.Hö., T.Ha., and Z.H. designed experiments. T.Ha. procured funding. C.B., N.A.P. and K.P. performed experiments. Z.H. supervised experimental work and analysis. T.Ha. and Z.H. wrote the manuscript with input from all co-authors. Correspondence and requests for materials should be addressed to T.H.

## Competing Interests

The authors declare no competing interests.

## Notes

### Competing Interest Statement

The authors have declared no competing interest.

